# Role of a kinesin motor in cancer cell mechanics

**DOI:** 10.1101/719054

**Authors:** Kalpana Mandal, Katarzyna Pogoda, Satabdi Nandi, Samuel Mathieu, Amal Kasri, François Radvanyi, Bruno Goud, Paul A Janmey, Jean-Baptiste Manneville

## Abstract

Molecular motors play important roles in force generation, migration and intracellular trafficking. Changes in specific motor activities are altered in numerous diseases. KIF20A, a motor protein of the kinesin-6 family, is overexpressed in bladder cancer, and KIF20A levels correlate negatively with the clinical outcome. We report here a new role for the KIF20A kinesin motor protein in intracellular mechanics. Using optical tweezers to probe intracellular mechanics and surface AFM to probe cortical mechanics, we first confirm that bladder cells soften with increasing cancer grade. We then show that inhibiting KIF20A makes the intracellular environment softer for both high and low grade bladder cancer cells. Upon inhibition of KIF20A cortical stiffness also decreases in lower grade cells, while it surprisingly increases in higher grade malignant cells. Changes in cortical stiffness correlate with the interaction of KIF20A with myosin IIA. Moreover, KIF20A negatively regulates bladder cancer cell motility irrespective of the underlying substrate stiffness. Our results reveal a central role for a microtubule motor in cell mechanics and migration in the context of bladder cancer.

## Introduction

Molecular motors convert chemical energy into mechanical work to achieve active cellular processes. The physical properties of molecular motors have been extensively studied *in vitro* by single molecular approaches ^1–3^. In cells, assemblies of molecular motors and collective active processes generate random fluctuating forces in the cytoplasm, which have recently been measured using force spectrum microscopy^4^. However, the individual contribution of a given type of molecular motor to cell mechanics is still unexplored. Kinesins form a superfamily of molecular motors that interact with microtubules and regulate key cell functions including mitosis, cell migration and organelle transport ^5–9^. We focus here on the role of the KIF20A kinesin (also called MKLP2 for mitotic kinesin-like protein 2 or Rabkinesin-6) in the mechanics of bladder cancer cells. KIF20A is a microtubule plus-end directed motor of the kinesin-6 family ^10^. KIF20A was initially shown to bind the small G protein RAB6 and to regulate retrograde transport from the Golgi apparatus to the endoplasmic reticulum (ER) during interphase ^11^. In mitosis, KIF20A localizes to the central spindle, and its phosphorylation is required for cytokinesis ^12^. A recent study shows that KIF20A and myosin II interact at Golgi hotspots to regulate the intracellular trafficking of RAB6-positive vesicles during interphase ^13^. KIF20A is involved in the fission of transport intermediates from the Golgi apparatus and serves to anchor RAB6 on Golgi and trans-Golgi network (TGN) membranes near microtubule nucleating sites ^11,13,14^. The regulation of myosin II might also be affected by KIF20A dynamics and thereby affect force generation and intracellular microrheological properties ^15–17^. It was recently reported that, during cortical neurogenesis, knockout of KIF20A causes the loss of neural progenitor cells and neurons due to early cell cycle exit and neuronal differentiation ^18^.

KIF20A was previously reported to be highly upregulated in many cancer cell types such as pancreatic cancer, melanoma, bladder cancer, liver cancer and breast cancer ^19–25^. Overexpression of KIF20A is associated with increased proliferation and tumor progression, poor prognosis and increased drug resistance in many cancers ^26,27^. In contrast, downregulation of KIF20A reduces cell proliferation in pancreatic cancer and glioma cells due to cytokinesis failure, appearance of binucleated cells, and apoptosis ^26,28^. However it has also been proposed that KIF20A could play antagonistic roles by both activating and inhibiting tumor progression ^19^. Peptides derived from KIF20A have been used to activate the immune system to kill cancer cells, which endogenously express the KIF20A antigen ^29^. Hence, KIF20A has become an immunotherapeutic target for several cancers, such as pancreatic or breast cancers ^29–31^.

Cell stiffness is often altered in pathological situations including fibrosis and cancer ^32–36^. In most cancers, isolated cancer cells have been shown to be softer than their healthy counterparts ^36,37^. Cancer cell softening is thought to be a key event during the metastatic process because more deformable cells should migrate more efficiently through confined environments to form metastases at a distant site from the primary tumor. Since actomyosin contractility has been shown to be crucial for maintaining cell stiffness and for force generation ^38,39^, a large body of research has been dedicated to study the role of actomyosin in cell mechanics ^40–42^. In contrast, the role of the microtubule cytoskeleton and its associated motors in cell stiffness has been much less studied. In particular, how microtubule motors participate to cell stiffness and how deregulation of kinesins in disease correlate with cell mechanics are still open questions.

In this study, we investigate the mechanical role of KIF20A in bladder cancer cells using a combination of active microrheological intracellular measurements by optical tweezers and atomic force microscopy (AFM) indentation experiments. We focus here on the role of KIF20A in regulating intracellular and cortical mechanics and correlate the mechanical effects of KIF20A with its effects on cell migration. Our results suggest that the interaction of KIF20A with myosin II differently regulates intracellular and cortical mechanics. The role of KIF20A in cell mechanics and cell migration points to KIF20A as a potential therapeutic target for bladder cancer.

## Results

### Bladder cancer cells are softer than normal urothelial cells

We first compared the viscoelastic properties of primary normal human urothelial (NHU) cells and two bladder cancer cell lines, grade II RT112 cells and grade III KU19-19 (KU) cells using intracellular active microrheology and AFM indentation experiments (Fig. 1). Cells were plated on crossbow shaped adhesive micropatterns to normalize their shape and intracellular organization. Internalized 2 µm diameter beads were displaced in an oscillatory fashion using optical tweezers-based rheology to measure the intracellular shear modulus (Fig. 1A, see Methods). We found that the intracellular shear modulus was almost twice higher in normal NHU cells than in bladder cancer cells and was significantly reduced in grade III cells compared to grade II cells (Fig. 1A). Intracellular relaxation experiments of beads trapped by optical tweezers (Fig.1B) allowed us to measure two phenomenological parameters, the rigidity index and the bead step amplitude, and to extract the complex shear modulus using a model based on power law rheology (see Methods). These measurements confirmed that lower grade (RT112) cells are softer than higher grade (T24) cells ^36,37^ (Fig. 1B). AFM indentation experiments showed that stiffness is also lower in grade III cells (Fig. 1C). Note that the stiffness measured by AFM indentation experiments was larger by two orders of magnitude than the intracellular stiffness measured either by oscillation or relaxation experiments, suggesting that AFM measurements are dominated by cortical stiffness. Stiffness maps obtained on micropatterned cells show a strong spatial dispersion of stiffness values (Fig. 1D) with higher values in the perinuclear region in both RT112 and KU cells, as observed using intracellular rheology ^43^. Despite the spatial variability of the elasticity measurements, the perinuclear region appears to be softer in grade III cells than in grade II cells (Fig. 1.D), consistent with the spatially averaged data (Fig. 1C).

**Figure 1:**
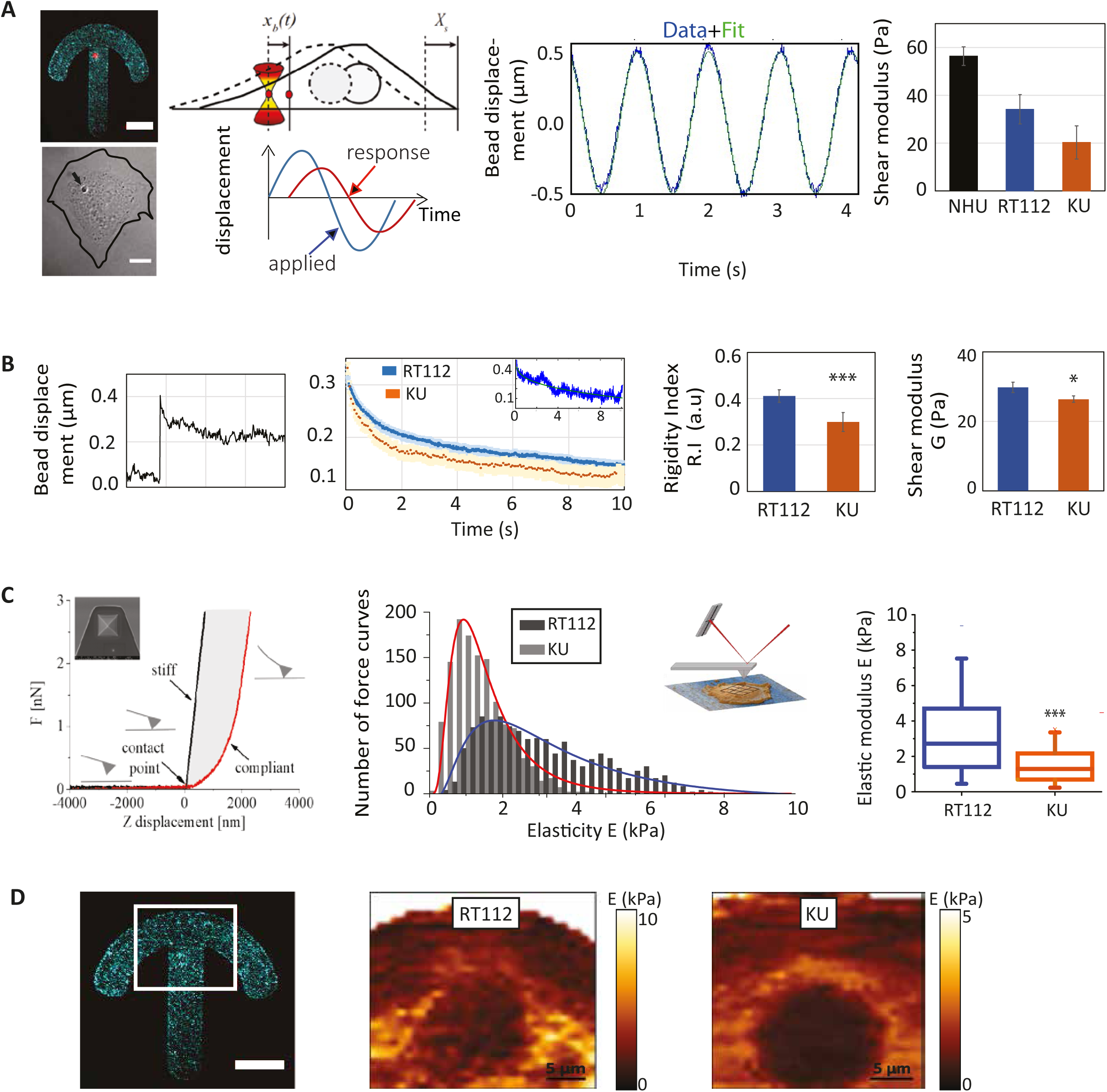
Bladder cancer cells are softer than normal urothelial cells and soften with increasing grade. (A) Intracellular oscillations experiments. (Left) Typical fluorescent and corresponding brightfield images of a normal human urothelial cell plated on a fibronectin-coated crossbow shape micropattern (cyan) with an internalized bead (red). The line shows the cell contour. Scale bars, 10 μm. (Middle) Cartoon showing the bead initially trapped at the center of the optical tweezers at time t =0 s, while the cell is moved in an oscillatory fashion. Typical oscillatory displacement of the bead and fitting of the curve (see Methods). (Right) Shear modulus measured by intracellular oscillating optical tweezers microrheology in normal urothelial (NHU) cells, grade II (RT112) and grade III (KU) bladder cancer cells. Data are from N=52, 20 and 20 cells for NHU, RT112 and KU cells respectively. Error bars represent standard errors. p-values are determined from Student’s t-test for unpaired samples (*** p<0.0001; * p<0.01). (B) Intracellular relaxation experiments. (Left) Relaxation curves showing the bead displacement from the center of the optical trap as a function of time following a .5 µm step displacement of the stage for RT112 cells (blue) and KU cells (orange). The relaxation strongly depends on the bead microenvironment: a stiff (blue) or soft (orange) microenvironment is characterized by a faster or slower relaxation respectively. The inset shows an example of curve fitting using a power-law model (see Methods). (Right) The bar graphs show the averaged rigidity index and shear modulus of RT112 and KU cells calculated from the relaxation curves using a phenomenological model (for the rigidity index) or a power-law model (for the shear modulus). Data are from N=36 and 60 cells for RT112 and KU cells respectively. Error bars represent standard errors. p-values are determined from Student’s t-test for unpaired samples (*** p<0.0001; * p<0.01). (C) AFM indentation experiments. (Left) Schematic representation of the AFM experiment. A quadratic pyramid geometry tip with an opening angle of 36° has been used for the experiment. Typical single point force-distance curve. Glass was used for calibration. The force increases after the contact point. Glass shows an infinite stiffness (black curve) while a cell is compliant (red curve). (middle) Distribution of the elasticity E for RT112 cells and KU cells. (Right) Averaged elasticity of RT112 cells and KU cells. Data are from N= 60, 60 cells for NHU and KU cells respectively. Error bars represent standard errors. p-values are determined from Student’s t-test for unpaired samples (*** p<0.0001; * p<0.01). (D) Spatial maps of the elasticity E measured on micropatterned cells. (Right) The red box shows the region from which the spatial maps were taken. Scale bar, 10 µm. (Center and left) Single cell stiffness maps of RT112 (center) and KU (right) cells. Scale bars, 5 µm.

### Inhibiting KIF20A softens the cell cytoplasm

Out-of-equilibrium active cellular processes impact strongly on cell rheology ^44^. While acto-myosin contractility is well-known to play a major role in the mechanical properties of cells and in force generation, the role of the microtubule-associated kinesin motors in cell mechanics is much less studied. Because the expression levels of several molecular motors including KIF20A have been shown to be deregulated in bladder cancer ^19^, we asked whether the kinesin KIF20A could participate in bladder cancer cell mechanics. We inhibited KIF20A by treating RT112 and KU bladder cancer cells plated on glass coverslip with 50 µM paprotrain, a specific KIF20A inhibitor, for 1 hr ^45^. Cells were treated with DMSO in the corresponding control experiments. Viscoelastic relaxation experiments showed an intracellular softening of both RT112 and KU cells upon inhibition of KIF20A (Fig. 2). The rigidity index, the bead step amplitude, and the storage modulus decrease significantly upon treatment with paprotrain for both cell lines, while the loss modulus decreases significantly only for RT112 cells (Fig. 2B-D).

**Figure 2:**
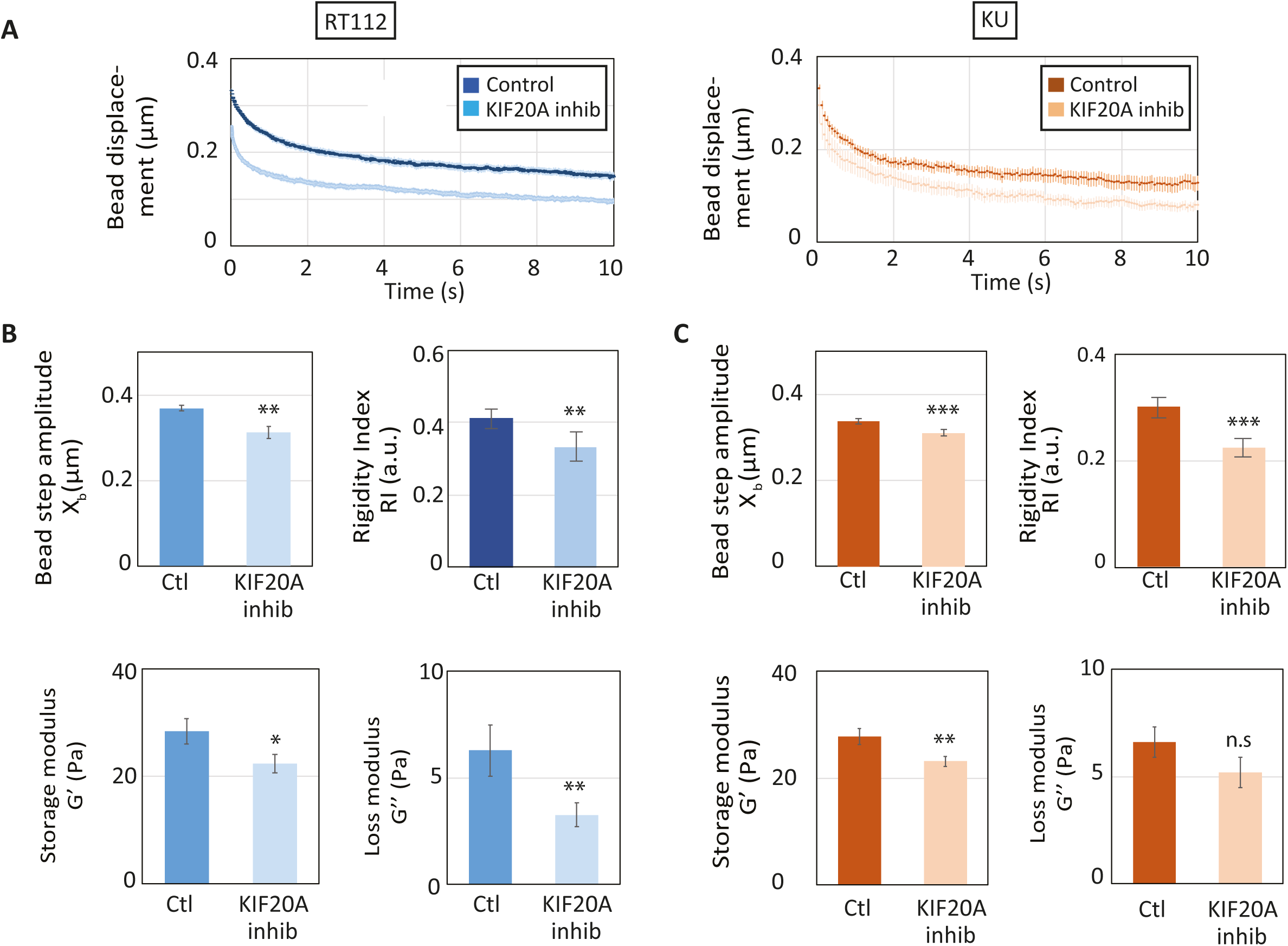
KIF20A inhibition softens the cytoplasm of both low and high grades bladder cancer cells. (A) Average intracellular relaxation curves for control RT112 (blue) and KU (orange) cells for control and for cells treated with the KIF20A inhibitor paprotrain (light blue, light orange). (B-C) Rigidity index, bead step displacement, storage modulus G’ and loss modulus G’’ in control and KIF20A inhibited RT112 cells (B) or KU cells (C). Data are averages from N=36 and 21 cells for control and KIF20A inhibited RT112 cells and from N=60 cells in both control and KIF20A inhibited KU cells. Error bars represent standard errors. P values are determined from Student’s t-test for unpaired samples with respect to control cells (*** p<0.001; ** p< 0.01; * p<0.05; and n.s. p>.05 not significant).

### KIF20A regulates cortical stiffness and cell motility on both soft and stiff substrates

To correlate intracellular rheology with the mechanical properties of the cell cortex, we measured cortical stiffness in RT112 and KU cells by AFM indentation experiments. Surprisingly, while grade II RT112 cells became softer when KIF20A was inhibited, grade III KU cells became stiffer (Fig. 3 A, B).

**Figure 3:**
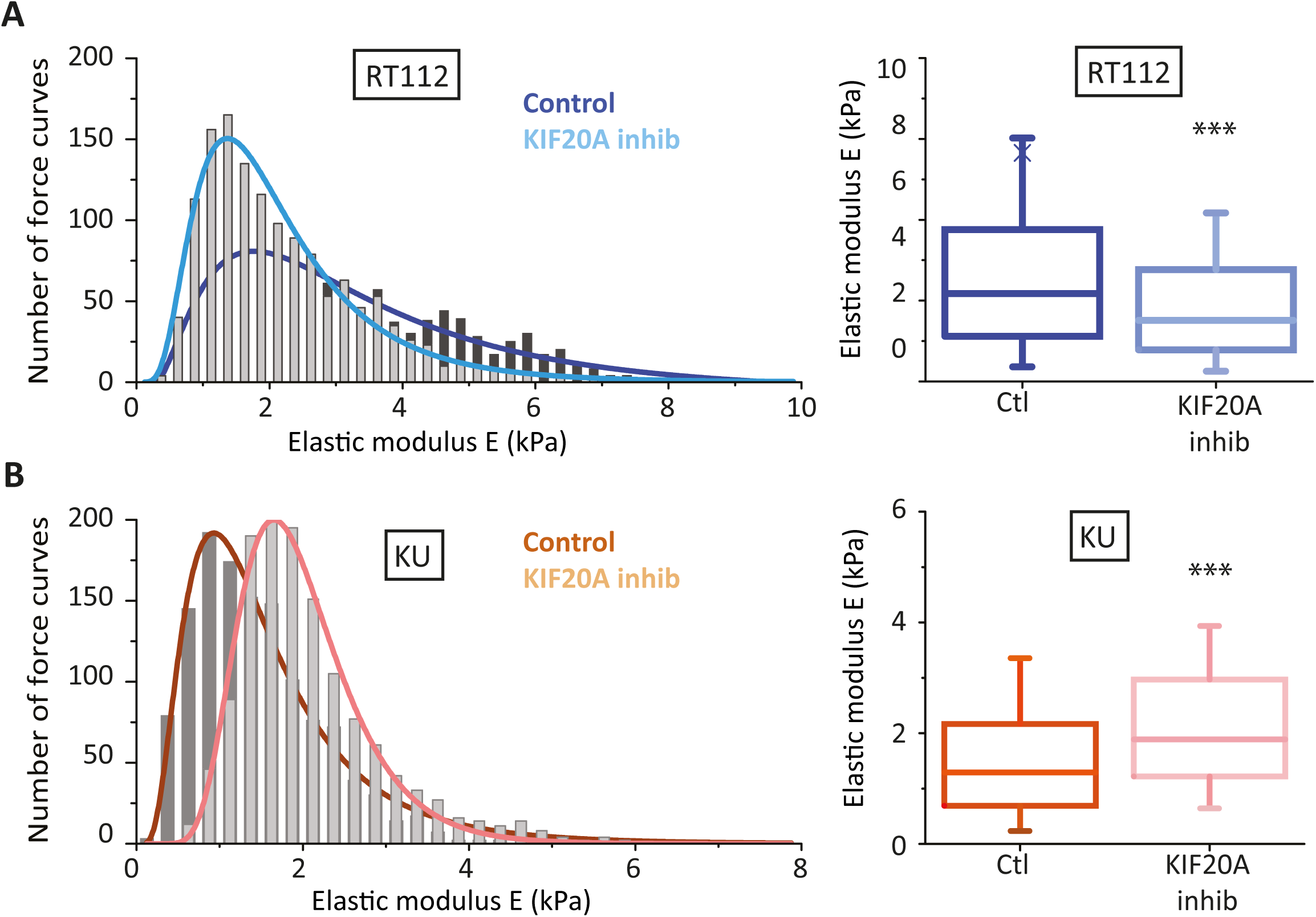
KIF20A inhibition softens the cortex of low-grade bladder cancer cells but stiffens the cortex of high grade bladder cancer cells. (A) (Left) Distribution of the elasticity E measured by AFM indentation experiments for control grade II RT112 cells (black) and for RT112 cells treated with the KIF20A inhibitor (grey). (Right) Averaged elasticity of control RT112 cells (blue) and RT112 cells treated with the KIF20A inhibitor (light blue). (B) Same as (A) for KU cells. Data are from N=60 cells per condition and for each cell about 25 force vs. distance curves were acquired. Distributions are fitted with a log normal distribution. Error bars represent standard errors. p values are determined from Student’s t test for unpaired samples with respect to control cells (*** p<0.0001; ** p<0.001; and * p<0.01).

The effects of KIF20A on bladder cancer cell rheology suggest that KIF20A impacts cellular functions that depend on cell mechanics such as cell migration. We studied individual 2D cell motility by plating RT112 and KU bladder cancer cells on soft (G=500 Pa) polyacrylamide hydrogels or on stiff (glass) substrates (Fig. 4). On both substrates, the higher-grade KU cells moved about twice as fast as the lower grade RT122 cells (Fig. 4 A, B and supplementary movies 1 and 2). When KIF20A was inhibited, the motility of RT112 and KU cells decreased. The speed of KU cells decreased about 3-fold and 4-fold on stiff and soft substrates respectively. In contrast, KIF20A inhibition had less effect on RT112 cell motility with only a 1.5-fold and 1.1-fold decrease in cell speed on stiff and soft substrates respectively (Fig. 4A, B and supplementary movies 3 and 4).

**Figure 4:**
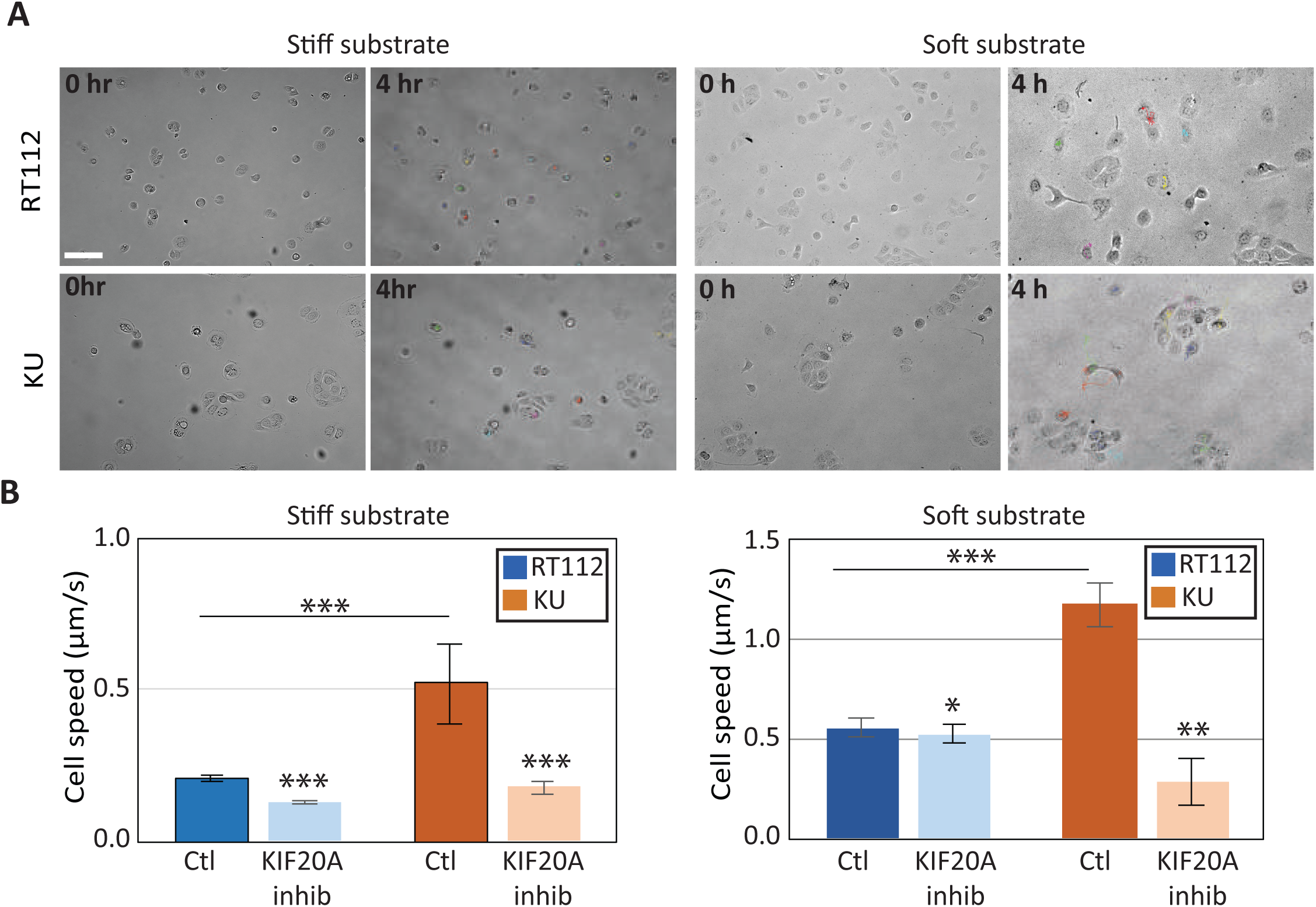
KIF20A inhibition reduces individual bladder cancer cell motility. (A) Bright-field images of randomly migrating RT112 cells (upper panels) and KU cells (lower panels) plated on stiff (glass, left) and soft (G=500 Pa polyacrylamide hydrogel, right) substrates at t=0 hr and t=4 hrs time points. Cell tracks are shown in color in the t=4hr images (see supplementary movies S1-S4. Scale bar, 100 µm. (B) Average cell speed measured from images acquired every 5 min for 5-8 hours Error bars represent standard error mean. On soft substrates, data are from N=25 and 22 control and KIF20A inhibited RT112 cells respectively, and N=24 and 18 control and KIF20A inhibited KU cells respectively. On stiff substrates, N=20 and 18 control and KIF20A inhibited RT112 cells respectively, and N=39 and 22 for control and KIF20A inhibited KU cells respectively. Error bars represent standard errors. p-values are determined from Student’s t-test for unpaired samples (*** p<0.0001; ** p<0.001, * p<0.01).

### Inhibiting KIF20A affects the subcellular localization of cortical actin and myosin II

We have shown so far that the effects of KIF20A inhibition are different in grade II RT112 cells and in grade III KU cells. First, KIF20A inhibition decreases cortical stiffness in RT112 cells, whereas it increases cortical stiffness in KU cells (Fig. 3). Second, KIF20A has stronger effects on single cell motility in KU cells than in RT112 cells, especially on soft substrates (Fig. 4). To explain these differences, we have first compared the expression levels and localization of KIF20A in both cell lines. RT112 cells express higher levels of KIF20A than KU cells (Fig. 5A). Since RT112 and KU have different cell cycle durations, we also measured the KIF20A expression levels in synchronized cells and found a similar trend as in non-synchronized cells, but with higher amounts of KIF20A (Fig. 5A).

**Figure 5:**
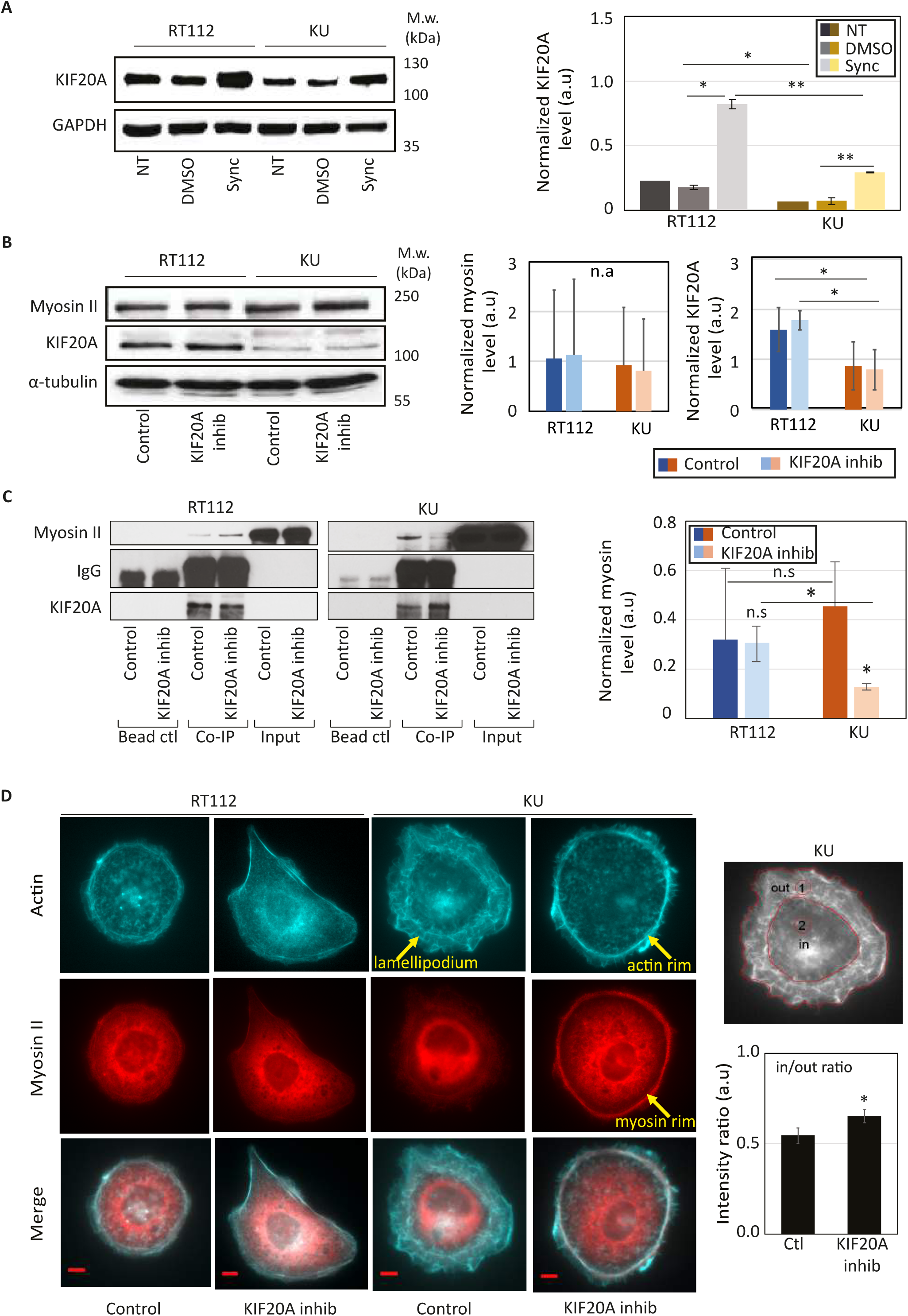
KIF20A interacts with myosin II in bladder cancer cells and its inhibition affects cortical acto-myosin organization specifically in high grade bladder cancer cells. (A) (Left) KIF20A expression in non-treated (NT), control (DMSO) or synchronized in mitosis (Sync) RT112 and KU cells (top). (Right) Quantification of the blots showing the normalized expression of KIF20A relative to the loading control (GAPDH). Error bars represent standard deviation of N=2 independent experiments. p-values are determined from Student’s t-test for unpaired samples (** p<0.005, * p<0.05, n.s p> 0.05 not significant). (B) (Left panel) Levels of myosin II and KIF20A in RT112 and KU cells treated with DMSO (control) or with 50 µM paprotrain (KIF20A inhibition). (Right panel) The graph shows quantification of the KIF20A and myosin band intensities normalized by the loading control (GAPDH or tubulin). Error bars represent standard deviation of N=3 independent experiments. (C) Interaction between myosin II and KIF20A shown by co-immunoprecipitation in control cells treated with DMSO and in cells treated with the KIF20A inhibitor (paprotrain 50 µM) in both RT112 and KU cells. (Left panels) Endogenous Myosin II is pulled down using a KIF20A antibody. Myosin II bound to KIF20A and KIF20A were revealed by western blot analysis using an anti-Myosin II antibody and an anti-KIF20A antibody respectively. Note that myosin II is detected in the input due to its high expression levels as opposed to KIF20A. (Right panel) Quantification of the myosin II/KIF20A interaction with respect to the total amount of KIF20A showing a reduced interaction after KIF20A inhibition in KU cells. Error bars represent standard deviation of N=3 independent experiments. p-values are determined from Student’s t-test for unpaired samples (** p<0.005, * p<0.05, n.s p> 0.05 not significant). (D) Actin and myosin II localization in bladder cancer cells. (left) Immunofluorescence images of actin (cyan, top), myosin II (red, middle) and merged actin/myosin II (bottom) for RT112 cells (right panels) and KU cells (left panels) treated with DMSO (control) or with the KIF20A inhibitor. Scale bars, 10 μm. (right) Myosin distribution was quantified by measuring the myosin fluorescence from an outer (lamellipodium, ‘1’) region and an inner (central, ‘2’) region (left). Normalized intensities plotted for control and KIF20A inhibited KU cells (right). Error bars represent standard errors (*N*=16 and 15 cells for control and KIF20A inhibited cells respectively). p-values are determined from Student’s t-test for unpaired samples (* p<0.05).

Because KIF20A interacts with myosin II ^13^, inhibition of KIF20A may have an effect on myosin II localization and/or cell contractility and hence cortical stiffness and cell motility. We first checked that KIF20A inhibition does not alter the expression levels of KIF20A and myosin II (Fig. 5B) and that the previously reported KIF20A/myosin II interaction also occurs in bladder cancer cells (Fig. 5C). The interaction between KIF20A and myosin II was comparable in RT112 and KU cells but was specifically reduced in KU cells when KIF20A was inhibited (Fig. 5C). We then asked whether KIF20A inhibition could affect KIF20A localization and the distribution of cytoskeletal fibers. While KIF20A inhibition did not affect the colocalization of KIF20A with microtubules and myosin in the perinuclear region (Fig. 5D and S1), the distribution of actin and myosin II was clearly affected in KU cells, but not in RT112 cells, with actin and myosin accumulating at the cell periphery specifically in KU cells (Fig. 5D and S1). Such enrichment in peripheral contractile acto-myosin upon KIF20A inhibition in KU cells could be responsible for the increase in cortical stiffness and the effects on migration observed specifically in this cell line.

## Discussion

We report here that the KIF20A kinesin motor plays a major role in bladder cancer cell mechanics. We found that a grade II (RT112) and a grade III (KU) bladder cancer lines are softer than normal urinary human (NHU) cells at the intracellular scale (Figure 1A), consistent with previously published results^36,37^. When KIF20A was inhibited, intracellular rigidity decreased in both cell lines (Fig. 2). While a cortical softening concomitant with intracellular softening occurred in RT112 cells, KU cells exhibited a stiffer cortex upon KIF20A inhibition (Fig. 3). To our knowledge, this is the first observation of an effect of a microtubule motor on cortical mechanics.

The mechanical effects of KIF20A are mediated by its interaction with myosin II and subsequent regulation of the subcellular localization of actin and myosin II. The KIF20A/myosin II interaction was originally described to occur through Rab6 at the Golgi apparatus^13^. We have focused here on the effects of KIF20A inhibition at the cell cortex, but it is likely that inhibition of KIF20A also perturbs post-Golgi trafficking and more generally intracellular transport. We propose that two pools of myosin II interact with KIF20A, a first pool localized at the Golgi apparatus and a second pool localized at the cell cortex (Fig. S1, S2). Inhibition of KIF20A may affect the Golgi pool in both RT112 and KU cells and reduce intracellular stiffness by softening internal membranes ^43^ such as Golgi membranes where KIF20A and myosin interact.

Additional effects of KIF20A inhibition on intracellular transport or microtubule dynamics, especially those of Golgi-nucleated microtubules, may also contribute to intracellular softening. In KU cells which display lower basal levels of KIF20A, the KIF20A/myosin II interaction is decreased when KIF20A is inhibited, which may release myosin II from Golgi membranes and allow enrichment of acto-myosin at the cortex. Such an increase in cortical acto-myosin density may lead to the stiffening of the cortex and to the reduced cell motility observed specifically in KU cells (Fig. 4). Because RT112 cells have a higher basal level of KIF20A and inhibition of KIF20A does not impact its interaction with myosin II in these cells (Fig. 5 and see discussion below), KIF20A inhibition could perturb the cortical pool of myosin II to a lesser extent and have a lower effect on cell motility in RT112 cells. Traction force measurements could help clarify the effects of KIF20A inhibition on cell-substrate adhesion and on cell motility^41,46,47^.

The different effects of KIF20A inhibition in RT112 and KU cells can likely be explained by the difference in KIF20A expression levels (Fig. 5A) and by the different strength of the previously reported KIF20A/myosin II interaction ^13^ (Fig. 5B-C). We found that inhibiting KIF20A did not affect the levels of KIF20A or myosin II in either cell line (Fig. 5B) but strongly reduced the KIF20A/myosin II interaction in KU cells but not in RT112 cells (Fig. 5C). In parallel with the reduced KIF20A/myosin II inhibition, actin and myosin reorganized at the cortex of KU cells (Fig. 5D and Fig. S2). A circular acto-myosin cortical rim appeared at the periphery of KU cells treated with the KIF20A inhibitor replacing the lamellipodial protrusions observed in control KU cells. This finding may explain both the loss of motility of individual KU cells and the increased cortical rigidity upon KIF20A inhibition. Similarly, KIF20A was reported to participate in actin rearrangement and protrusion formation in pancreatic cancer cells ^48^. In contrast no significant change in cortical acto-myosin organization was observed in RT112 cells, which correlates with the lower effects of KIF20A inhibition on RT112 cell motility.

Because microtubules play a central role in cell division, they have been early targets of cancer drug treatments. For instance, taxol (Paclitaxel) blocks mitosis by stabilizing microtubules and is a widely used chemotherapy drug. Similarly, microtubule-associated molecular motors are potential targets for cancer therapy ^49^. Kinesins and dynein play a crucial role in many cellular functions such as mitosis and cell proliferation and, not surprisingly, the expression levels of kinesins are often modified during tumor progression. Here we have used the specific KIF20A inhibitor paprotrain to study the effects of KIF20A inhibition on mechanical phenotypes in RT112 and KU bladder cancer cells. Our results point to KIF20A as a potential target for bladder cancer therapy. KIF20A inhibition had stronger effects on single cell motility in KU cells than in RT112 cells (Fig. 4). The velocity decreased by a factor greater than two in the case of KU cells on stiff or soft substrates (Fig. 4B). Thus, KIF20A is involved in two hallmarks of malignant transformation, hyper-proliferation and, as we show here through its role in cell mechanics, increased cell motility. We can thus speculate that a KIF20A inhibitor could be of considerable therapeutic interest as it would not only target cancer cell proliferation but also motility and probably invasion.

## Material and Methods

### Cell culture and reagents

RT112 and KU19-19 (KU) cells were grown in RPMI medium supplemented with 10% (vol/vol) FBS and 1% Penicillin/Streptomycin at 37 °C with 5% (vol/vol) CO_2_. Primary normal human urothelium (NHU) (Southgate et al., 1994) cells were grown in KSFMC medium (Life Technologies) supplemented with 5% (vol/vol) horse serum. For intracellular rheology, cells were incubated overnight with 2-μm-diameter fluorescently labeled latex beads (660/690 fluorescence; Bangs Laboratories) before seeding them on the micropatterns. To inhibit KIF20A, cells were incubated for at least 1 hr with 50 μM Paprotrain (Biokinesis, gift from Stéphanie Miserey-Lenkei) before intracellular rheology and AFM measurements. 10µM Hepes was added before microrheology and AFM experiments. For immunofluorescence experiments, cells were fixed with 4% (vol/vol) paraformaldehyde. Primary antibodies were anti- β tubulin (Sigma; T4026), anti-myosin, and phalloidin (Invitrogen), DAPI (Sigma), Myosin II (Sigma), KIF20A (Tocris bioscience). Secondary antibodies were from Jackson Immuno Research Laboratories, E-cadherin (Cell Signaling), vinculin (Santa Cruz).

### Adhesive micropatterns

Crossbow-shaped micropatterns were printed on PEG-coated glass coverslips by deep UV photolithography, and then coated with 50 μg/mL fibronectin and 20 μg/mL Alexa 546–fibrinogen-red (Sigma). Cells were seeded on freshly prepared protein-coated micropatterns and allowed to spread for at least 2–3 hrs before experiment started ^50^. Non-adherent cells were washed off by rinsing with culture medium.

### Optical tweezers-based intracellular microrheology, image acquisition, and data processing

The setup combining optical tweezers and fast confocal microscopy was described in detail previously ^43^. Briefly, a single fixed optical trap was built on an inverted Eclipse microscope (Nikon) equipped with a resonant laser confocal A1R scanner (Nikon), a 37 °C incubator, and a nanometric piezo stage (Mad City Labs). Coverslips with micropatterned or non-patterned cells containing typically one to three internalized beads were mounted in a Ludin chamber or glass-bottom dishes (MatTek Corporation) were used. Trapped beads were subjected either to an automated oscillatory displacement of 0.5-μm amplitude (intracellular oscillations experiments) or to a 0.5 µm step displacement (intracellular relaxation experiments). Images were recorded at 120 or 15 frames per second in the resonant mode of the confocal scanner for 40 s using the NIS Nikon software. The bead position was tracked using a homemade Matlab single particle tracking routine ^43^.

#### Intracellular relaxation experiments

We assume that the creep function *J(t)* behaves as a power law with an exponent *α* and a prefactor *A, J(t)* = *At*^*α*^ as described in^43^ (see methods). The bead position *x*_*b*_(*t*) was fitted using:

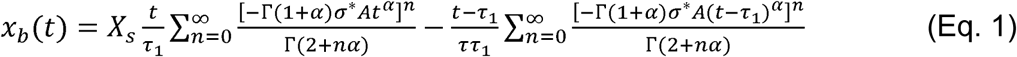

We use the first four terms of Eq.1 to fit the first ten seconds of the relaxation of the bead position *x*_*b*_(*t*) as a function of time knowing the stage step displacement *x*_*s*_ = 0.5 μ*m* and the duration of the step *τ*_1_ = 40 ms (the stage step displacement is modelled by a linear increase from 0 μ*m* to *x*_*s*_ = 0.5 μ*m* between *t* = 0 and *t* = τ_1_ seconds), and 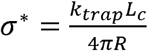 is a geometrical factor involving the trap stiffness *k*_*trap*_ = 210 pN, the bead radius *R* = 1 μ*m* and a characteristic length *L*_*c*_.

The characteristic length *L*_*c*_ was chosen so that the generalized Stokes relation is verified^17,51,–53^: *K* = 6*πRG*, where *G* is the shear modulus and *K* the apparent complex stiffness of the bead microenvironment. In our case, *G* = *σ/ε*, where 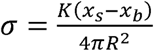 is the applied stress with *x*_*b*_ and *x*_*s*_ the bead and the stage displacements respectively, and *ε* = (*x*_*s*_ − *x*_*b*_)/*L*_*c*_ is the strain of the microenvironment. Thus, *G* = *KL*_*c*_ /(4*πR*^2^), and taking into account the generalized Stokes relation, *G* = *K*/6*πR*, we find *L*_*c*_ = 2*R*/3 = 0.67 µm and σ^*^ = 12.7 ±0.5 Pa.

The prefactor *A* and the exponent *α* are the only two fit parameters in Eq. 1. Because bead tracking was noisier at longer timescales due to intracellular dynamics, the data was fitted using the following weights: 0.9, 0.09 and 0.01 for 0<t<2s, 2<t<5s and 5<t<10s respectively. The complex shear modulus *G* = *G*’ + i*G’’* was deduced from *A* and *α* using:

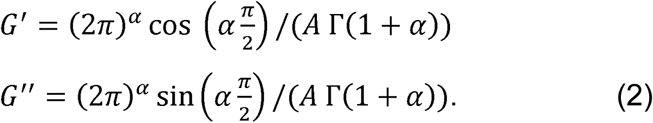

As in ^43^, we also used a phenomenological index to quantify intracellular rigidity which we called the ‘rigidity index’ (*RI*) and defined as: 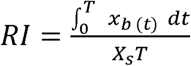. *RI* varies between 0 and 1. The bead step amplitude *X*_*b*_ corresponds to the initial displacement of the bead following the step displacement of the stage (*X*_*s*_=0.5 µm > *X*_*b*_).

#### Intracellular oscillations experiments

The bead is initially trapped at the center of the optical tweezers at time t =0 s. An oscillating displacement of the stage *x*_*s*_(*t*) = *x*_*s*_ sin(*ωt*) was applied at a frequency *ω*=1 Hz for 10 sec and amplitude *x*_*s*_ = 0.5 μm ^54^, with *x*_*s*_ <*x*_*s*_ and Δ*θ* the pase lag due to the visoelastic natutre of thecytoplasm.

The force applied on the bead trapped in the optical tweezers of stiffness *k*_*trap*_ = 214 pN,is 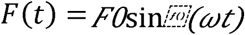, with *F0=ktrapxs* and the corresponding bead displacement is *xbt=xbsin ωt − Δ θ* where sthe phase lag due to the viscoelastic nature of the, where the phase lag Δ*θ* is 0 for a purely elastic material and *π*/2 for a purely viscous material. The complex shear modulus is given by

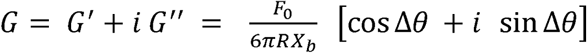

Where,

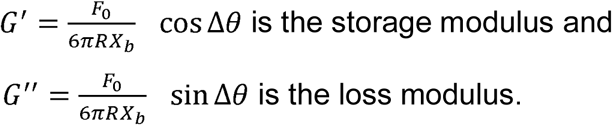

The frequency was fixed to *ω*=1 Hz and the amplitude of the stage displacement was *x*_*s*_ = 0.5 μ*m*. The amplitude of the bead displacement *x*_*b*_ and the phase lag Δ*θ* were determined using sinusoidal fitting with a homemade Matlab routine after tracking the bead position as described above.

### AFM based rheology, image acquisition, and data processing

Atomic force microscopy (AFM)-based microindentation was performed using a Dimension Icon AFM (ORC Bruker Nano) and a quadratic pyramid tip (Opening angle = 36°) with stiffness value k = 0.1N/m. At each indentation location, the probe tip was programmed to indent the cell at a 15 μm/s constant z-piezo displacement rate (equal rate of retraction) if the indentation depth 0.5 μm with force value 2 nN maximum indentation force. The applied force was 2 nN for a 500 µm indentation depth at an approach and retraction velocity of 15 µm/s. Glass was used for calibration. For each cell, indentation was performed on relatively flat regions (avoiding cell edges) of 5 μm × 5 μm of cell surface area in contact mode surface scans. Force-distance indentation curves are collected from 25 different locations for each cell. For each indentation curve, the cantilever deflection (in volts) and z-piezo displacement (in μm) were converted to an indentation force (in nN) and depth (in μm) through calibrating the cantilever deflection sensitivity (nm/V) by indenting on a hard mica substrate and a spring constant (nN/nm) via thermal vibration. The loading portion of the curve at each location was fitted to the elastic Hertz model via least-squares linear regression to calculate the effective indentation modulus at the given indentation rate^55^. To determine the elasticity of the cell every individual curve is fitted by the Hertz model. Whole cell stiffness is determined by the width of stiffness (E) distributions when fitted with a log normal distribution (Fig3B). Force 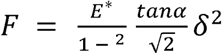; where E* is the effective Young’s modulus, 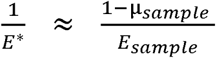; = Poisson’s ratio, α = face angle; δ = indentation depth

### Hydrogel preparation

For single cell random migration, cells were cultured either on glass (stiff substrate) or 500 Pa polyacrylamide hydrogels (soft substrate). 500 Pa polyacrylamide hydrogels were prepared by mixing 7 % acrylamide and 0.28 % bis-acrylamide in water. APTMS and TEMED were added to initiate gel polymerization. A sulfo-SANPAH solution in water was added and the gel surface was activated by UV light. Gels were coated with 100 mg/ml collagen type I.

### Single cell migration

RT112 and KU cells were plated at a density of 30,000 cells per gel or on bare glass 22×22 mm coverslips for 24 hrs at 37 °C with 5% (vol/vol) CO_2_. Cells were treated with DMSO (control) or 50 μM paprotrain before starting time-lapse acquisition. 5 hrs to 8 hrs time lapse sequences are acquired with a frame every 5mins. Image J manual tracking plugin is used to find the cell displacement and velocity. Velocities between two time points are obtained and averaged over total time period.

### Western Blot

KU and RT112 cells were either non treated, treated with DMSO or treated with Nocodazole (0.4 mg/ml) for 18h to synchronize the cells. Cells were lysed in a buffer containing 150 mM NaCl, 50 mM Tris-HCl pH 7.5, 1% Nonidet-P40 (Sigma). Protein concentrations were determined by Quick Start™ Bradford 1x Dye Reagent (Bio-Rad). For Western blot analysis a nitrocellulose Protran BA 83 membrane (Life science) was used and the signal from HRP-conjugated secondary antibodies was detected with an ECL system and an enhanced chemiluminescence system (ChemiDoc Touch System, Bio-Rad). Quantification of the mean band intensity relative to the loading control Glyceraldehyde 3-phosphate dehydrogenase (GAPDH) was performed using the Image Lab software.

### Co-Immunoprecipitation

The previously reported interaction between myosin II and KIF20A was tested by co-immunoprecipitation in bladder cancer cells. RT112 and KU cells were treated with 2 mg/ml nocodazole for 16 hrs. Cells were washed and supplemented with fresh media to recover. Cells were either treated with DMSO or with 50 μ M paprotrain for 1 hr, washed twice in PBS, and incubated on ice for 60 min in a lysis buffer (25mM Tris pH 7.5, 50 mM NaCl, 0.1% NP40, with freshly added protease and phosphatase inhibitor cocktails (Sigma). Cells were clarified by centrifuging at 10,000g for 10 minutes. 500 µg of cell lysates were processed for co-immunoprecipitation using 5 µg of KIF20A antibody coupled to Protein A-dyna beads overnight at 4 °C in lysis buffer. Bead control was performed using 500ug of cell lysate without KIF20A antibody. Beads were washed four times in 1ml of lysis buffer followed by heating in 2X Laemmli sample buffer. The immunoprecipitates were analyzed by SDS-PAGE followed by Western immunoblotting. For western-blotting experiments, cell solubilization was performed in lysis buffer. The following primary antibodies were used: rabbit anti-MHC (Covance; 1:2000), rabbit anti-KIF20A (Bethyl or A17425; 1:1000). Secondary horseradish peroxidase (HRP)-coupled antibodies were from Jackson Laboratories.

### Statistical analysis

Data are expressed as means ± standard error mean. Statistical relevance was evaluated using Student’s t-tests and the p-value is indicated (n.s no significant; * p<0.05, ** p<0.01, *** p<0.001 unless otherwise stated).

## Acknowledgements

We thank Stephanie Miserey-Lenkei for useful the gift of Paprotrain and technical assistance of KIF20A western blots. We thank Atef Asnacios and Charlotte Alibert for technical support and discussions. KM was supported by a grant from MERSEC, Physical Science Oncology Center (PSOC) grant U54-CA193417 and Labex postdoctoral fellowship. KP was funded Physical Science Oncology Center (PSOC) grant U54-CA193417. SM was funded by a grant from Sorbonne Université UPMC University Paris 06 (Programme Doctoral “Interfaces Pour le Vivant”). AK was funded by a grant from the Agence Nationale de la Recherche (ANR). PAJ acknowledges NIH grant GM111942. This work was also supported by the Center for Engineering Mechanobiology through NSF STC 1548571. JBM acknowledges funding from INSERM Plan Cancer 2009-2013 INSERM - CEA Tecsan (grant number PC201125).

